# Patched regulates cell cycle and tissue architecture in *C. elegans* gonad

**DOI:** 10.1101/2025.06.09.657693

**Authors:** Johanna Farley, Madeleine Schwalbe, Fredrik Forsberg, Aqilah Amran, Sandeep Gopal

## Abstract

Hedgehog (Hh) signaling is essential for embryonic development and tissue homeostasis in several organisms. However*, Caenorhabditis elegans* lacks canonical Hh signaling due to the absence of key components, such as smoothened (SMO) and Hh ligands. Despite this, *C. elegans* retains Patched homologs, *ptc-1* and *ptc-3*, which have specialized independent functions. Although *ptc-1* is predominantly expressed in the gonads and *ptc-3* in somatic tissues, we demonstrate that both genes are required to maintain germ cell populations and proper actin cytoskeletal architecture in the progenitor zone of the germline. Disruption of actin-encoding genes impairs germ cell cycle progression and reduces germ cell numbers, indicating that cytoskeletal integrity is critical for maintaining the germline. Furthermore, defects observed upon loss of Patched function are linked to disruptions in cholesterol metabolism. We show that the phenotypes observed in the gonads due to the loss of Patched function can be rescued by modulating dietary cholesterol. Together, our findings reveal a role for Patched receptors in regulating germline architecture and germ cell development through cholesterol-sensitive functions, offering insights into how metabolic cues influence the organization of complex tissues.

## INTRODUCTION

Hedgehog (Hh) signaling is critical for embryonic development, tissue homeostasis, and stem cell maintenance(1, 2). The canonical hedgehog pathway is ligand-dependent and requires Hh proteins, such as Sonic Hedgehog, Indian Hedgehog or Desert Hedgehog, to initiate signaling. In the absence of Hh proteins, the transmembrane receptor Patched (PTCH) inhibits the activity of Smoothened (SMO), a G protein-coupled receptor-like protein, to block Hh signaling. PTCH relieves SMO inhibition upon engagement with Hh proteins, likely through direct transport of cholesterol(3–5). Active SMO initiates a transcriptional signaling cascade that regulates cell proliferation, differentiation, and survival. Aberrant functioning of Hh signaling pathways contributes to metabolic disorders, tumor growth, metastasis, and drug resistance(1, 2, 6, 7).

Unlike vertebrates, which have a well-defined Hh signaling pathway involving Hh ligands, PTCH, SMO, and associated transcription factors, *Caenorhabditis elegans* lacks homologs of several core components of the canonical Hh pathway(8). However, it possesses hedgehog-related genes and proteins that regulate important developmental processes(8, 9). *C. elegans* expresses two PTCH orthologs, PTC-1 and PTC-3(8, 10). Early loss of *ptc-1* or *ptc-3* leads to embryonic lethality, suggesting their critical functions during development(11, 12). While *ptc-1* appears to be predominantly expressed in the gonads, *ptc-3* is expressed in somatic tissues with very low expression in the gonad(13, 14). Despite the absence of SMO and other canonical pathway components, *ptc-1* and *ptc-3* are involved in important signaling pathways in *Caenorhabditis elegans*. For instance, *ptc-1* is essential for cytokinesis, whereas *ptc-3* is important for cholesterol transport and lipid storage in *C. elegans*(15, 16). This study aimed to elucidate the specific roles of *ptc-3* and *ptc-3* in the regulation of germ cell proliferation and gonad architecture in *C. elegans*, with a particular focus on the impact of cholesterol intake on these processes. *C. elegans*, with its well-characterized *ptc* orthologs, offers a unique model to dissect the role of *ptc-1*and *ptc-3* in non-canonical signaling pathways affecting germ cell behavior.

Adult hermaphrodite *C. elegans* carry two syncytial germline tubes, within which the germ cell nuclei are partially enclosed by membranes(17–19). Germ cell development begins distally within the progenitor zone, where a small number of self-renewing germline stem cells produce new germ cells(20). Germ cell proliferation is under combinatorial control by multiple intrinsic and extrinsic signaling pathways and environmental cues(20–24). Germ cells in adult animals undergo proximal differentiation to produce mature oocytes(25, 26). The gonad maintains a specific architecture supported by an actin cytoskeleton that assumes characteristic structure at each part of the gonad(24, 27, 28). Previous reports have revealed that the actin structural pattern is critical for determining germ cell behavior and protein distribution in the gonad(24, 27).

Here, we report the critical functions of Patched in the regulation of germ cell proliferation and gonad architecture. Specifically, we found that *ptc-1* and *ptc-3* are involved in maintenance of germ cell cycle and gonad architecture in *C. elegans*. We also show that cholesterol intake by the animals is critical for maintaining gonad architecture, and the defects caused by the loss of Hh receptors can be rescued by modifying cholesterol intake by the animals. This highlights the importance of environmental and metabolic cues in rectifying germline defects and fertility in animals.

## MATERIALS AND METHODS

### *C. elegans* maintenance

Wild-type hermaphrodite *C. elegans* strains were maintained on Nematode Growth Medium plates seeded with 400 μL of OP50 *Escherichia coli* at 20 °C, unless otherwise stated.

### RNA interference (RNAi) plasmid preparation

The sequences of *the ptc-1* and *ptc-3* genes were amplified using the forward and reverse primers, and HindIII sites were introduced at both ends of the sequence. The amplified insert and empty vector (L4440) were digested using HindIII and ligated together using DNA ligase. The plasmid sequences were verified using Sanger sequencing and transformed into HT115(DE3) E. coli. RNAi plasmids for *act-3* and *act-4* were obtained from the Ahringer RNAi library, and their sequences were verified by Sanger sequencing.

### RNAi experiments

HT115 *E. coli* bacteria expressing RNAi plasmids for *ptc-1, ptc-3*, *pos-1* genes and an empty vector (L4440), were grown in Luria Broth (LB) + Ampicillin (100 μg/mL) (Sigma #A9518) at 37 °C, 220 RPM for 16 h. 400 μL of RNAi bacterial cultures were spread on NGM plates supplemented with 1 mM IPTG (Merck #420291) and 50 µg/mL carbenicillin and dried for at least 48 h at room temperature in the dark. L4 stage worms were fed with RNAi bacteria for 20 h. To confirm the efficacy of the RNAi plates and protocol, animals silenced for the *pos-1* gene, which causes embryonic lethality, were monitored for 36 h.

### Brood size analysis

L4 hermaphrodites were picked onto individual RNAi plates seeded with HT115 *E. coli* bacteria expressing RNAi plasmids for *ptc-1, ptc-3, pos-1* genes or an empty vector. Worms were allowed to lay eggs for 24 h, and the mothers were then moved to new individual RNAi plates. After another 24 h, embryos from the first plate were analyzed for hatching. This process was repeated for 5 days. The number of larvae and embryos was counted each day and summed as the brood size.

### Germline analysis

Following RNAi, worms were placed in a 10 µL droplet of 0.01% levamisole on a coverslip for sedation. Germ lines were extruded by dissecting the sedated worms. The coverslips containing the germ lines were then inverted onto poly L-lysine-coated slides and transferred to liquid nitrogen for 1 min. The coverslip was then gently but quickly separated from the glass slide to expose the adhered germline. The slides were immediately fixed in a freshly prepared fixing solution containing 0.08 M HEPES (pH 6.9), 1.6 mM MgSO_4_, 0.8 mM EGTA, and 3.7% PFA in PBS for 25 min at room temperature. The slides were washed for 5 min in phosphate-buffered saline (PBS, pH 7.4) containing 0.1% Tween-20 (PBST). The fixed germ lines were permeabilized for 10 min in PBS containing 0.2% Triton-X and washed twice with PBST for 5 min. Germ lines were then blocked using 30% normal goat serum for 20 min, and the excess blocking solution was removed using a paper towel. Germ lines were incubated with 1:10,000 4ʹ,6-diamidino-2-phenylindole (DAPI) and 1:4000 phalloidin-555 in NGS for 1 h at room temperature in the dark. After staining, the germ lines were washed twice with 0.1% PBST for 5 and 10 min, respectively. Slides were mounted by applying a drop of Fluoroshield mounting media on the germ lines, followed by a coverslip and dried at 4°C until imaging. Germ lines were imaged using the Leica Stellaris 5 Confocal Laser-Scanning Inverted Microscope with a 63x objective, and the cell number in the progenitor zone was analyzed using Imaris software, as described previously(28). The distance between three nearest nuclei were measured using a built-in tool of Imaris(24). Distance graphs were plotted by normalizing the values against the control RNAi. For phalloidin staining phenotypic analysis, the percentage of germ lines with actin cytoskeletal structures deviated from the control germlines were calculated. Phalloidin staining intensity after actin gene silencing was calculated using the formula, Corrected Total Phalloidin Intensity (fluorescence) = Integrated Density − (area × mean gray background). The graphs were plotted by normalizing the values against the control. To accommodate the likely changes in the total area of measurement after actin RNAi, the values were divided by the area of the progenitor zone. For S-phase analysis, after RNAi, the germlines were labelled using 5-ethynyl-2ʹ-deoxyuridine (EdU) by soaking worms in 250 μM EdU solution for 15 min(24). The germlines were then extruded and stained using the Click-iT Alexa Flour (488) EdU labeling kit (Invitrogen), followed by 2 h of DAPI staining. The S-phase stages were identified and quantified based on previously reported analysis(24, 29).

### Cholesterol experiments

To study the effect of cholesterol on germ cell development, L4 hermaphrodite worms were placed on plates containing 0, 5, or 20 mg/mL cholesterol, while animals were fed with bacteria containing RNAi plasmids. For each condition, worms were fed bacteria containing RNAi plasmids for *ptc-1, ptc-3*, or the control for 20 h. Young adult worms were collected and dissected, and the germlines were analyzed as described above.

### Lipid staining

RNAi NGM plates were prepared as described above, with the addition of 1 mL of 1 M filtered glucose or 1 mL mMQ water as a control. The plates were seeded with bacteria containing RNAi plasmids for *ptc-1, ptc-3*, or the control. 30 L4 worms were placed on the RNAi plates at 20°C overnight. For lipid staining, Oil Red O stock (ORO) solution was prepared (0.5% w/v) in isopropanol and mixed at room temperature for 2 days in the dark. Two hours prior to staining, the ORO stock solution was diluted to 60% with sterile mQ water and filtered through a 0.22 µm filter. The worms were collected and washed with PBS to remove the bacteria. The worms were fixed in 3 parts fixation solution (30 mM PIPES, 14 mM Na_4_EGTA, 40 mM NaCl, 160 mM KCl, 0.4 mM spermine, 4% formalin, 0.5% beta-mercaptoethanol) diluted in 1 part PBS. Tubes containing worms were immersed in ethanol and dry ice bath for 3 min, followed by thawing for 1-2 min. This cycle was repeated twice. The worms were washed three times in 1X PBS prior to adding 60% isopropanol and incubating at room temperature on a rotating rack for 10 min. The worms were incubated with 60% ORO solution, completely protected from light, and incubated for 4-6 hours at room temperature on a rotating rack. Stained worms were washed twice with 0.01% PBST and once with 1X PBS. Worms were transferred to a 2% agarose pad on a glass slide, and a coverslip was gently placed over the worms. Worms were imaged immediately on an Olympus BX61VS brightfield Slide Scanner at 20x objective with the proximal intestinal cells in focus. Images were analyzed using Fiji (ImageJ 2.14.0/1.54f). For each worm, approximately the first 4 sets of intestinal cells (after the pharynx and prior to the helical twist) were selected for measurement. Area, mean gray value and integrated density were measured, and a background measurement taken from an unstained area inside the head. To calculate the corrected total lipid intensity based on fluorescence for the selected area in each worm, the following formula was used: Corrected Total Lipid Intensity (fluorescence) = Integrated Density − (area × mean gray background). The graphs were plotted as normalized values against control RNAi.

### Statistical Analysis

Ordinary one-way ANOVA with Tukey’s multiple comparison test was performed to compare the groups. Samples from at least 3 independent experiments were analyzed (Refer source data). Statistical significance was set at p < 0.05.

## RESULTS

### Patched receptors control the cells in the progenitor zone

Short-term silencing of Patched receptors, *ptc-1* and *ptc-3,* from late L4 stage resulted in a reduced number of progenies produced by the hermaphrodite *C. elegans* (Fig. 1A). This suggests an aberrant germline function. To test the impact of Patched in the germ line, we silenced *ptc-1* and *ptc-3* from the late L4 stage, when gonad development nears its completion. We found that the number of germ cell nuclei in the progenitor zone was reduced in both *ptc-1* and *ptc-3* silenced animals, suggesting that germ cell production was negatively affected (Fig. 1B-C). It has been shown that germ cells proliferate and differentiate in the folded gonad tissue(27, 28). These structures are supported by a spiral-shaped actin cytoskeleton (Supplementary fig. S1)(27, 28). To test whether the germline architecture is affected by Patched receptors, we stained the germline cytoskeleton using phalloidin and analyzed the cytoskeletal features in the germline using confocal microscopy (Fig. 1D, Supplementary fig S1). We found that the germline cytoskeleton was significantly altered after silencing *of ptc-1* and *ptc-3* (Fig. 1D-E, Supplementary fig. S1). The tissue folds maintained by cytoskeletal support can determine the distribution of nuclei within the progenitor zone, which can potentially determine the access of nuclei to proteins expressed in the distal germline (24, 27). Therefore, we analyzed the nuclear distribution after silencing Patched in the germ line (Fig 1F). This revealed that the relative distance between the nuclei in the progenitor zone is significantly reduced in the gene silenced animals in comparison with control animals (Fig. 1F-G). Taken together, our results indicate that Patched controls the germline architecture and cell number in the progenitor zone of the germ line.

**Figure 1.**
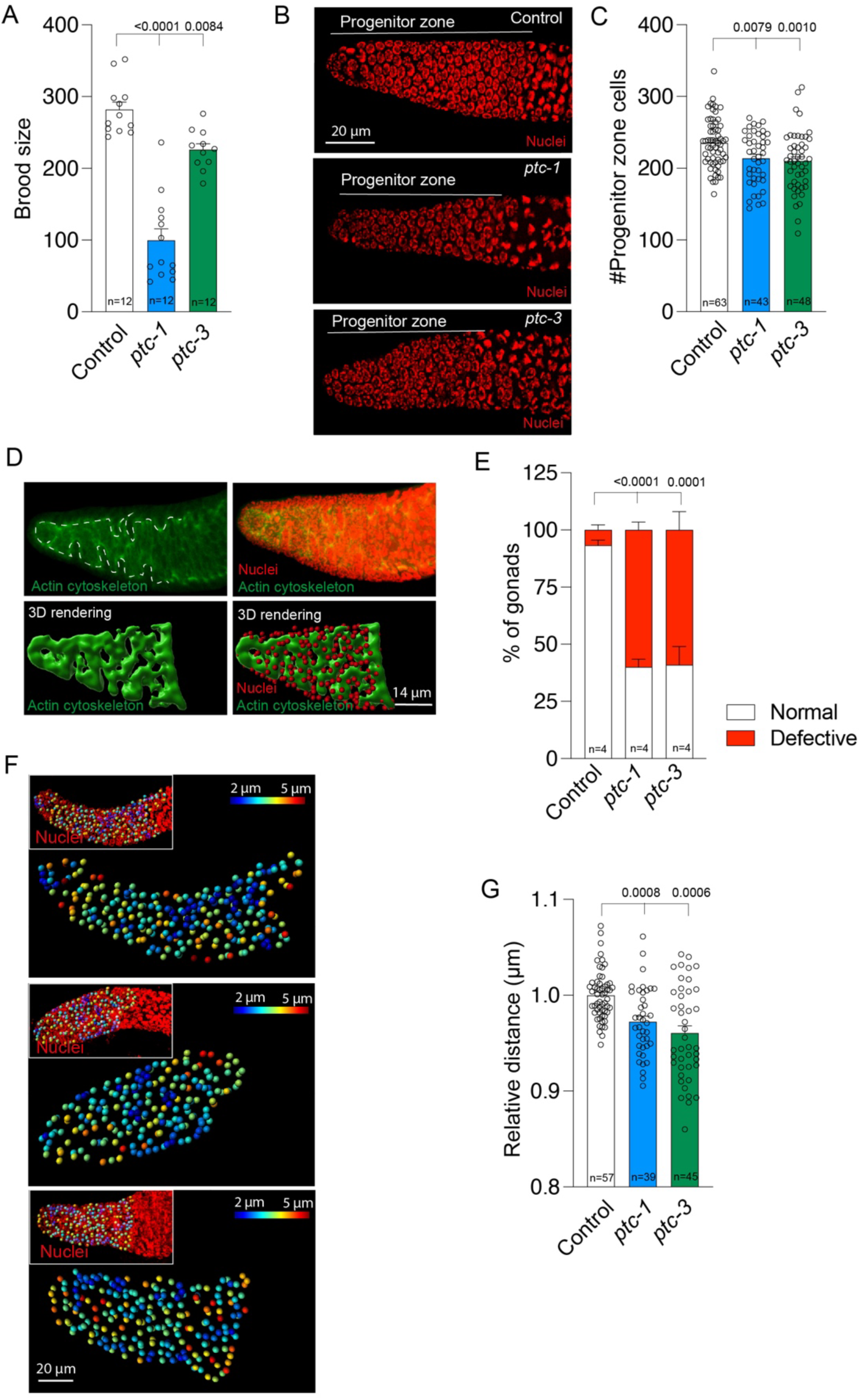
Patched receptors control the distal germline. A) The loss of *ptc-1* or *ptc-3* from late L4 to one day old adult animals results in a reduced brood size. B) Images of distal gonad in control, ptc-1 and ptc-3 RNAi. DAPI (red) nuclei staining of distal germline. Progenitor zone marked in the images. Scale bar, 20 µm. Quantification of the number of cells at the progenitor zone in control and *ptc-1* or *ptc-3* silenced animals. D) Phalloidin (actin cytoskeleton - green) and DAPI (nuclei - red) staining showing the typical actin morphology of distal gonad. Supplementary fig. S1. Top panel – Confocal images projected to 2D surface. Bottom panel – 3D rendering of actin cytoskeleton and nuclei in the progenitor zone. Scale bar, 14 µm. E) Quantification of percentage of gonads with normal and defective cytoskeleton in control and *ptc-1* or *ptc-3* silenced animals. Experiment was performed in 4 replicates with a total of 70 germlines for control, 108 germlines for *ptc-1* RNAi and 78 germlines for *ptc-3* RNAi. F) Distribution of nuclei within the progenitor zone in control and *ptc-1* or *ptc-3* silenced animals. 3D rendering shows the placement of nuclei. Each nucleus is color coded based on the average distance with its closest three neighboring nuclei. Insets show the 3D rendering of the nuclei placed on projected confocal images. Scale bar, 20 µm. G) Quantification of the relative distance of each nucleus with its closest 3 neighboring nuclei in *ptc-1* or *ptc-3* silenced animals in comparison to control animals. P values were assessed by one-way analysis of variance (ANOVA) with Tukey’s multiple comparison test. Error bars indicate SEM. P value < 0.05 = statistically significant.

### The actin cytoskeleton regulates germ cell behavior in the distal gonad

As the loss of *ptc-1* and *ptc-3* alters the gonad cytoskeleton and germ cell number at the distal gonad, we tested whether the cytoskeleton itself affects the germ cells. To study the impact of actin on germ cell behavior, we silenced *act-3* and *act-4* in late L4 worms and quantified the number of cells in the progenitor zone (Fig 2A, Supplementary Fig S3). Both actin isoforms are expressed in the gonads of *C. elegans* (Supplementary fig. S2)(14). The loss of *act-3* and *act-4* resulted in reduced total intensity and intensity per area of phalloidin staining suggesting critical loss of actin in the gonad (Fig. 2A-C). In addition, cytoskeletal structures showed defects in the progenitor zone after actin loss (Supplementary fig. S3). Silencing of both *act-3* and *act-4* resulted in a reduced number of cells in the progenitor zone (Fig. 2D). While a complete loss of the cytoskeleton in the germline after silencing either of the actins was not always observed within the gene knockdown window, the actin cytoskeleton appeared to be different from that in the control and the reduction in the expression was sufficient to induce changes in the number of cells in the progenitor zone (Fig. 2A-D, Supplementary fig. S3). The reduction in cell number could indicate a defective cell proliferation and cell cycle in the progenitor zone. To study the cell cycle, we analyzed the DNA synthesis (S phase) of the germ cells in the progenitor zone using 5-ethynyl-2ʹ-deoxyuridine (EdU) staining (Fig 2E, Supplementary fig. S4)(29). This revealed that silencing of both actins resulted in a reduced number of EdU-positive cells in the progenitor zone, suggesting a reduction in the number of cells in the S phase undergoing DNA synthesis (Fig 2E-I). However, *act-3* and *act-4* impacted the S phase to different extents. Silencing *act-3* affected the early and late S phases, whereas silencing *act-4* only affected the late S phase (Fig. 2G, 2I). The loss of both genes did not result in a significant change in the mid S phase (Fig. 2H). Collectively, our data show that actin-associated germ cell architecture is critical for maintaining germ cells in the progenitor zone.

**Figure 2.**
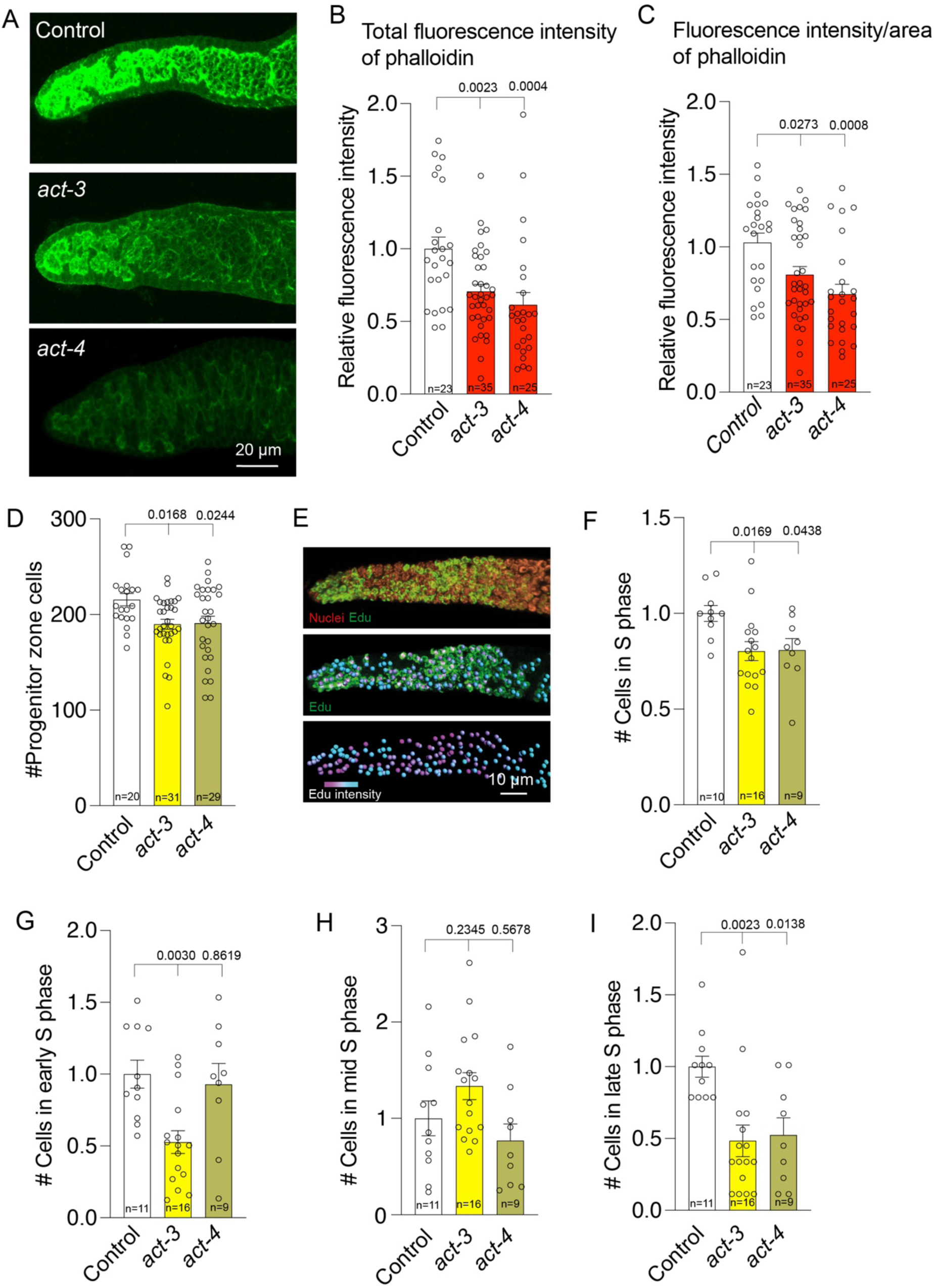
Actin cytoskeleton structure determines the cell behavior at the progenitor zone. A) Phalloidin staining showing the actin morphology of distal gonad in control and *act-3* or *act-4* silenced animals. Supplementary fig. S2, S3. B) Quantification total fluorescence intensity of phalloidin staining in the progenitor zone in control and *act-3* or *act-4* silenced animals. C) Quantification of phalloidin Fluorescence intensity per area of the progenitor zone in control and *act-3* or *act-4* silenced animals. D) Quantification of the number of cells in the progenitor zone in control and *act-3* or *act-4* silenced animals. E) EdU staining of the distal gonad. Top and middle panel-DAPI (nuclei - red) and EdU (S phase – Green) Bottom panel – 3D rendering of EdU positive nuclei with heatmap of EdU intensity. S phase stages were determined using Edu intensity and shape of the staining. Supplementary fig. S4. Scale bar, 10 µm. F) Quantification of the relative number of cells in S phase (all EdU positive) in *act-3* or *act-4* silenced animals in comparison to control animals. G-I) Quantification of the relative number of cells in early (G), mid (H) and late (I) S phase in *act-3* or *act-4* silenced animals in comparison to control animals. P values were assessed by one-way analysis of variance (ANOVA) with Tukey’s multiple comparison test. Error bars indicate SEM. P value < 0.05 = statistically significant.

### Cholesterol management is critical for determining germ cell fate in the progenitor zone

We hypothesized that if the actin cytoskeleton is disturbed by the loss of Patched, it would lead to an impaired cell cycle. To test this, we quantified the number of cells in the S phase of the cell cycle after silencing *ptc-1* and *ptc-3* (Fig. 3A-D). We found that the total number of cells in the S phase at the distal end was reduced after silencing both *ptc-1* and *ptc-3* (Fig. 3A)(29). We also found that silencing *ptc-1* significantly reduced the number of cells in the early and late S phases, whereas silencing *ptc-3* only reduced the number of cells in the late S phase (Fig. 3B, D). However, silencing of either of the Patched receptors did not result in a significant quantifiable difference in the mid S phase (Fig 3C). Previous reports suggested that Patched is critical in lipid management in *C. elegans*(*15*). The germ line in *C. elegans* is highly susceptible to nutrient intake and metabolism. The germline is reduced to a handful of stem cells in the absence of nutrients. First, we tested whether nutrient intake was affected by the loss of Patched receptors. We quantified the rate of pharyngeal pumping, which is essential for food intake, after silencing *ptc-1* and *ptc-3 genes.* We did not find a significant difference in pharyngeal pumping after the loss of *ptc-1* and *ptc-3*, which ruled out defective food intake as the reason for the reduced number of cells in the progenitor zone (Fig. 3E).

**Figure 3.**
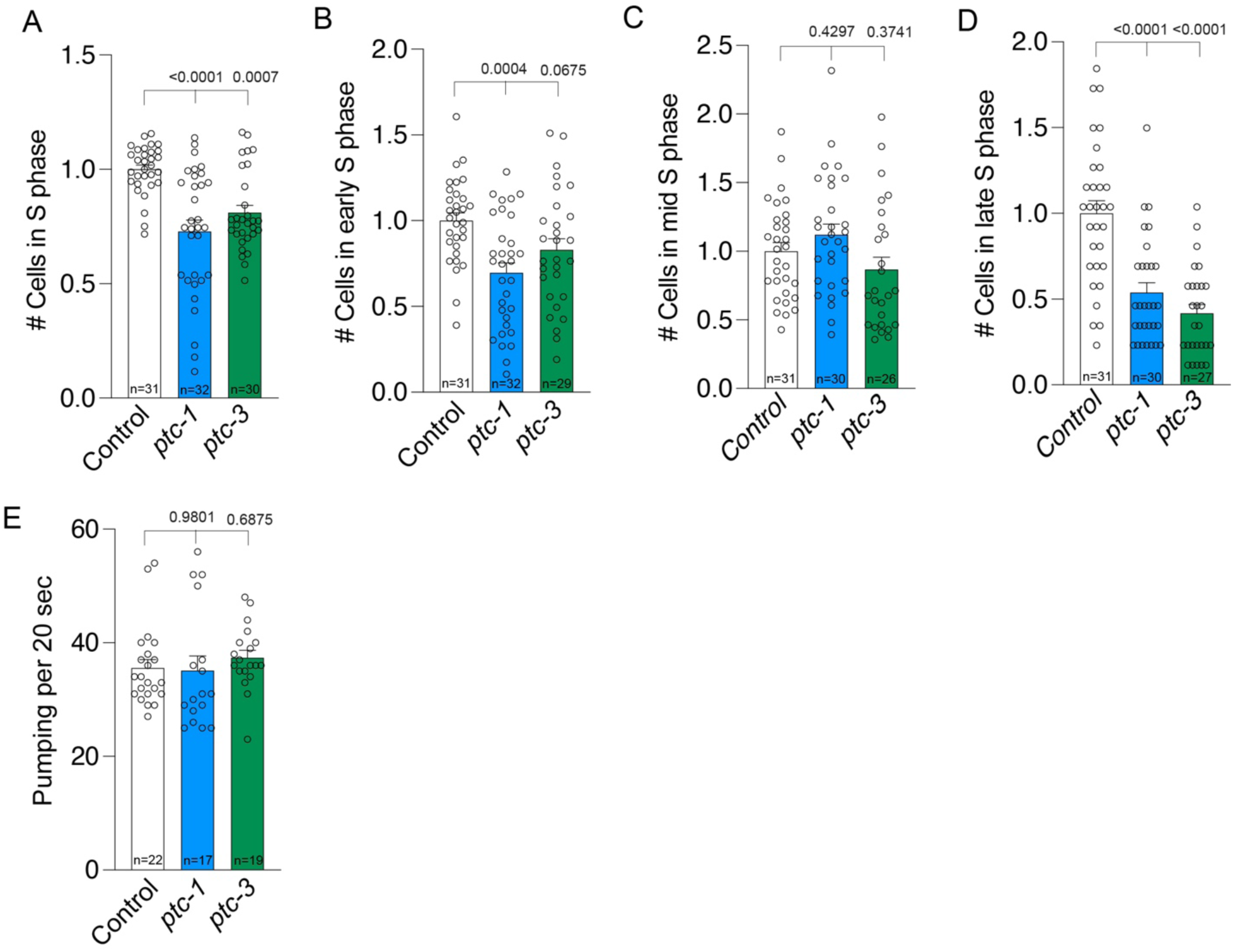
Patched regulates S phase of the germ cells in the progenitor zone. A) Quantification of the relative number of cells in S phase in *ptc-1* or *ptc-3* silenced animals in comparison to control animals. B-Quantification of the relative number of cells in early (B), mid (C) and late (D) S phase in *ptc-1* or *ptc-3* silenced animals in comparison to control animals. E) Quantification of pharyngeal pumping in control, *ptc-1* or *ptc-3* RNAi animals. P values were assessed by one-way analysis of variance (ANOVA) with Tukey’s multiple comparison test. Error bars indicate SEM. P value < 0.05 = statistically significant. P value < 0.05 = statistically significant.

Previous reports have shown that silencing *ptc-3* for an extended period (from the L2 stage) results in a reduction in total lipid storage in *C. elegans*(*15*). *C. elegans* stores the majority of lipids in the intestine. To test whether the loss of Patched receptors results in reduced intestinal lipid storage, we stained the animals with Oil Red after silencing *ptc-1* and *-3* from the late L4 stage for 20 h. We did not observe a reduction in fat levels in our experiments during the gene silencing period (Fig. 4A-B), suggesting that short-term silencing of Patched is not sufficient to produce a global impact on lipid storage in the animal. Furthermore, the increase in lipid levels due to a high glucose diet was not prevented by silencing either *ptc-1* or *ptc-3* (Fig. 4A-B)(15).

**Figure 4.**
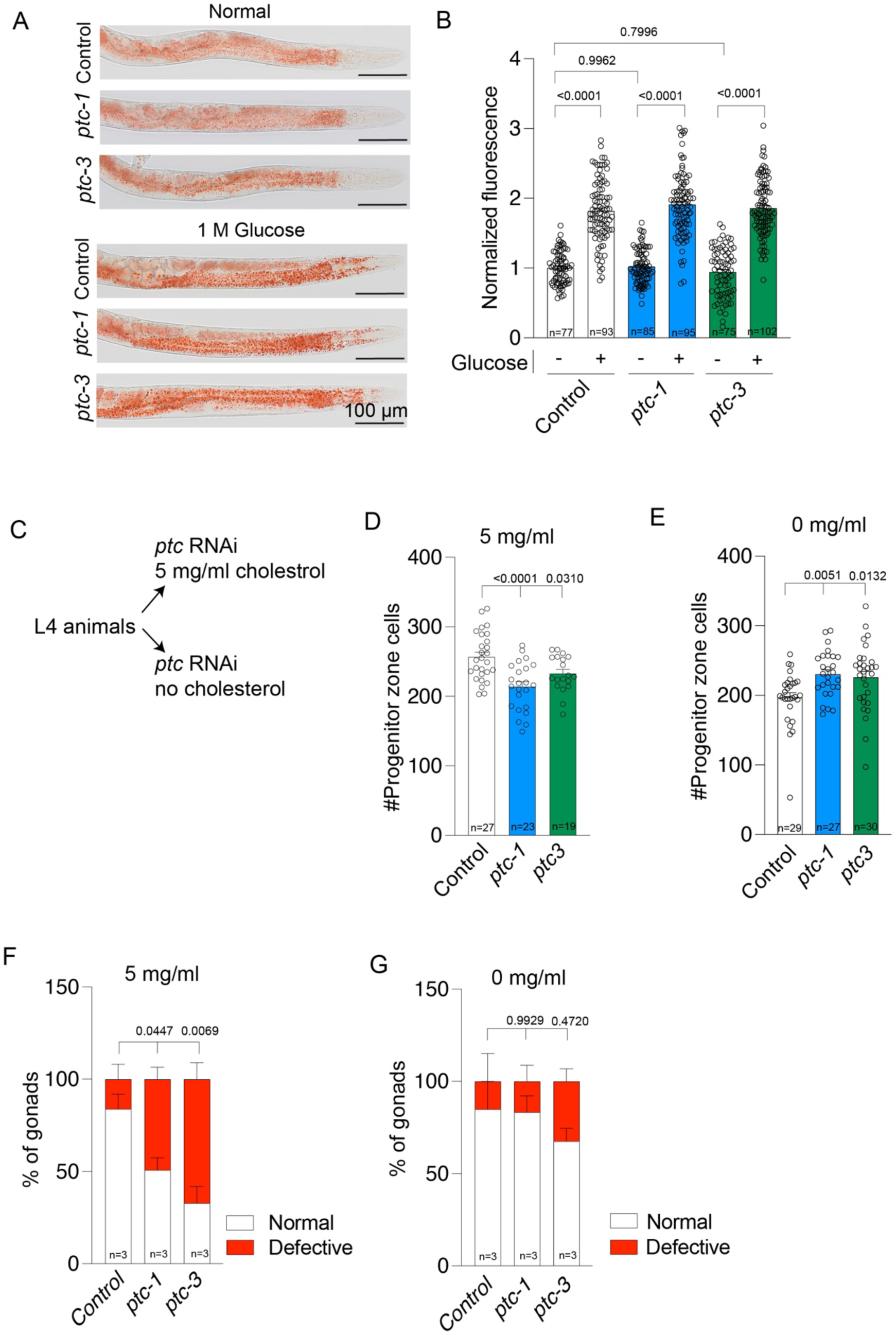
Patched control germ cells through cholesterol. A) Micrographs of ORO staining showing lipid content in animals under normal and high glucose (1M) diet. Worms placed on a high glucose diet marked with a visible increase in ORO intensity (red). Scale bar, 100 µm. B) Quantification of lipid content in control and Patched silenced animals under normal and 1M glucose. The total staining intensity was quantified in the intestine up to the helical twist and normalized against control values. C) Cholesterol experiment. L4 worms were placed on plates containing 5 mg/ml (normal) or no cholesterol while fed with bacteria containing RNAi plasmids for control, *ptc-1* and *ptc-3.* D) Quantification of the number of cells in the progenitor zone in animals grown under normal cholesterol conditions. Animals were silenced for control, *ptc-1* or *ptc-3*. Supplementary fig. S5, S6. E) Quantification of the number of cells in the progenitor zone in animals grown under no cholesterol conditions with RNAi. Animals were silenced for control, *ptc-1* or *ptc-3*. Supplementary fig. S5, S6. F) Quantification of percentage of gonads with normal and defective actin cytoskeleton in control, *ptc-1* or *ptc-3* silenced animals under normal cholesterol conditions. Experiment was performed in 3 replicates with a total of 25 germlines for control, 23 germlines for *ptc-1* RNAi and 14 germlines for *ptc-3* RNAi. G) Quantification of percentage of gonads with normal and defective actin cytoskeleton in control, *ptc-1* or *ptc-3* silenced animals under no cholesterol conditions. Experiment was performed in 3 replicates with a total of 21 germlines for control, 19 germlines for *ptc-1* RNAi and 27 germlines for *ptc-3* RNAi. P values were assessed by one-way analysis of variance (ANOVA) with Tukey’s multiple comparison test. Error bars indicate SEM. P value < 0.05 = statistically significant. P value < 0.05 = statistically significant.

Recent findings suggest that Patched acts as a cholesterol transporter and is critical for cholesterol maintenance in the cells(15). We hypothesized that the loss of *ptc-1* and *ptc-3* may have resulted in increased cholesterol accumulation in the cells. To test the impact of dietary cholesterol on the germ line, we fed the animals with varying levels of cholesterol. We found that changes in cholesterol levels in the NGM plates alone can impact the germ cell number in the progenitor zone (Supplementary fig. S5). A diet with no cholesterol (0 mg/ml) and excess cholesterol (20 mg/ml) resulted in a reduced number of cells in the progenitor zone compared to animals supplied with normal cholesterol levels (5 mg/ml) (Supplementary fig. S5). A report showed that the loss of Patched resulted in defective cholesterol efflux in cells(15). As the loss of Patched would have resulted in an increase in cellular cholesterol, we tested the impact of Patched silencing when animals were fed with no dietary cholesterol (Fig 4C-E). As expected, silencing *ptc-1* and *ptc-3* in animals supplied with 5 mg/ml cholesterol resulted in a reduced number of cells in the progenitor zone (Fig. 4D). We also found that the reduction in the number of cells in the progenitor zone due to a diet lacking cholesterol was rescued with a significant increase in cell number after silencing of *ptc-1* and *ptc-3* (Fig. 4D). In contrast, the reduction caused by excess cholesterol was not rescued by silencing *ptc-1* or *ptc-3* (Supplementary fig. S6). Next, we analyzed the effects of diet on gonad architecture. The cytoskeletal defects caused by silencing *ptc-1* and *ptc-3* were significantly reduced when the worms were on a diet lacking cholesterol (Fig 4F-G). These results collectively suggest that lowering dietary cholesterol can reduce the impact of cholesterol efflux defects caused by the loss of Patched function in animals and restore gonad architecture and germ behavior.

## DISCUSSION

Our study showed the role of two Patched homologs, *ptc-1* and *ptc-3*, in cell behavior at the progenitor zone of the *C. elegans* germ line. We found that both *ptc-1* and *ptc-3* are required to maintain brood size. This indicates that both receptors have important functions in the germline and germ cell development. Unsurprisingly, loss of *ptc-1* showed a stronger reduction in brood size, as it is predominantly expressed throughout the gonad and is expected to have more germline-specific functions. On the other hand, we found both *ptc-1* and *ptc-3* are required to maintain the number of cells in the progenitor zone, as the loss of either of the receptors caused significant reduction in the number of cells in the progenitor zone. Given the very low expression in the germline, it is surprising that *ptc-3* has a similar impact on the germline as *ptc-1*. It is plausible that *ptc-3* acts from other tissues and the impact is part of a global response in the animal(13, 14). The second defect we observed in the distal germ line was the impaired structure of the actin cytoskeleton. Actin organization is important for cytokinesis and the cell cycle(30, 31). Therefore, changes in actin structure may be relevant to germ cell proliferation. Reports suggest a role for *ptc-1* in cytokinesis of mitotically dividing cells based on the presence of multinucleated cells in the proximal germline, although there were no clear connections between *ptc-3* and germ cell behavior(16). Our analysis of the cell cycle (S phase/DNA synthesis) after silencing of Patched receptors confirmed a reduced number of cells in specific stages of the S phase, linking both *ptc-1* and *ptc-3* to the cell cycle. Previous reports have shown that germ cells proliferate and differentiate in folded tissue, which is supported by actin structures. It has been suggested that the nuclei and protein distribution within the progenitor zone could be dictated by the folded tissue architecture(27, 28). It is possible that changes in actin can disrupt this architecture and impact germ cell behavior. However, one can argue that the change in the actin cytoskeleton could be forced by over or under crowding of cells in the progenitor zone. While the actin organizational structure and cell cycle changed in the germ line after Patched silencing, it does not necessarily confirm that the actin changes can cause the germ cell changes. We addressed this prospect by silencing the actins expressed in the gonad. Our analysis of the germ line after silencing actin encoding genes *act-3* or *act-4,* two actins expressed in the gonad, revealed changes in S phase cells and a reduction in the number of cells in the progenitor zone. This suggests that the changes in actin isoforms indeed can directly impact the germ cells.

Our results showed that *ptc-3*, which has extremely low or no expression in the germline(14), also impacts germ cell behavior in the mitotic region. This suggests that the global functions of Patched receptors are also likely to affect germ cells. Recent studies showed that Patched receptors are key in managing lipid storage and cholesterol transport(5, 15, 32). In contrast to published data, our analysis did not show a reduction in lipid storage after silencing either *ptc-1* or *ptc-3* in *C. elegans*(*15*). Also, the silencing of these receptors failed to prevent glucose induced increase in lipid storage in the animals. It is likely that the silencing of Patched receptors, which was performed for a shorter time compared to published results, was not enough to impact the global lipid level in the animals. Further, we found that placing animals on low or very high cholesterol diets directly impacts the germline. In *C. elegans*, cholesterol acts as a membrane component and a precursor for hormones, though it is currently unclear how downstream signaling is impacted in the germline actin structure(33, 34). We found silencing Patched on a low cholesterol diet recovers the germ cell number and cytoskeletal defects at the progenitor zone. In addition, it is likely that changes in cholesterol levels impact membrane properties including stiffness, thus impacting the cell cycle progression(15, 35). Nevertheless, our data reiterates the role of Patched receptors in cholesterol efflux and their link to the germ line. Given the lack of a canonical hedgehog signaling pathway in *C. elegans,* it appears that Patched silencing led to unwanted cholesterol accumulation that affects the gonad and germ cell behavior in *C. elegans*. This shows an additional layer of control of germ cell development in *C. elegans*.

## DATA AVAILABILITY

Reagents and further information will be available upon request from the Gopal laboratory.

## ACKNOWLEDGMENTS

Some strains were provided by the *Caenorhabditis* Genetics Center (University of Minnesota), which is funded by the NIH Office of Research Infrastructure Programs (P40 OD010440). The gene family classification and functions were performed as curated in the Wormbase. Imaging for this project was performed at Lund Bioimaging Center, and Lund Stem Cell Centre Imaging Facility, which is supported by the Swedish national Stem Therapy initiative.

## DISCLAIMERS

Not applicable

## AUTHOR CONTRIBUTIONS

J.F., A.A and S.G. developed the concept. J.F., M.S., F.F., A.A. and S.G. developed the methodology. J.F., M.S., F.F. and A.A. performed the investigation. S.G. acquired funding. S.G. wrote the original manuscript. J.F., M.S., F.F. and A.A. performed review & editing.

## REFERENCES

1. Jing J, Wu Z, Wang J, Luo G, Lin H, Fan Y, et al. Hedgehog signaling in tissue homeostasis, cancers, and targeted therapies. Signal Transduct Target Ther. 2023;8(1):315.

2. Briscoe J, Therond PP. The mechanisms of Hedgehog signalling and its roles in development and disease. Nat Rev Mol Cell Biol. 2013;14(7):416–29.

3. Huang P, Nedelcu D, Watanabe M, Jao C, Kim Y, Liu J, et al. Cellular Cholesterol Directly Activates Smoothened in Hedgehog Signaling. Cell. 2016;166(5):1176–87 e14.

4. Teperino R, Aberger F, Esterbauer H, Riobo N, Pospisilik JA. Canonical and non-canonical Hedgehog signalling and the control of metabolism. Semin Cell Dev Biol. 2014;33:81–92.

5. Myers BR, Neahring L, Zhang Y, Roberts KJ, Beachy PA. Rapid, direct activity assays for Smoothened reveal Hedgehog pathway regulation by membrane cholesterol and extracellular sodium. Proc Natl Acad Sci U S A. 2017;114(52):E11141–E50.

6. Garg C, Khan H, Kaur A, Singh TG, Sharma VK, Singh SK. Therapeutic implications of sonic hedgehog pathway in metabolic disorders: Novel target for effective treatment. Pharmacol Res. 2022;179:106194.

7. Teperino R, Amann S, Bayer M, McGee SL, Loipetzberger A, Connor T, et al. Hedgehog partial agonism drives Warburg-like metabolism in muscle and brown fat. Cell. 2012;151(2):414–26.

8. Burglin TR, Kuwabara PE. Homologs of the Hh signalling network in C. elegans. WormBook. 2006:1–14.

9. Serra ND, Darwin CB, Sundaram MV. Caenorhabditis elegans Hedgehog-related proteins are tissue- and substructure-specific components of the cuticle and precuticle. Genetics. 2024;227(4).

10. Zugasti O, Rajan J, Kuwabara PE. The function and expansion of the Patched- and Hedgehog-related homologs in C. elegans. Genome Res. 2005;15(10):1402–10.

11. Sonnichsen B, Koski LB, Walsh A, Marschall P, Neumann B, Brehm M, et al. Full-genome RNAi profiling of early embryogenesis in Caenorhabditis elegans. Nature. 2005;434(7032):462-9.

12. Simmer F, Moorman C, van der Linden AM, Kuijk E, van den Berghe PV, Kamath RS, et al. Genome-wide RNAi of C. elegans using the hypersensitive rrf-3 strain reveals novel gene functions. PLoS Biol. 2003;1(1):E12.

13. Diag A, Schilling M, Klironomos F, Ayoub S, Rajewsky N. Spatiotemporal m(i)RNA Architecture and 3’ UTR Regulation in the C. elegans Germline. Dev Cell. 2018;47(6):785–800 e8.

14. Tzur YB, Winter E, Gao J, Hashimshony T, Yanai I, Colaiacovo MP. Spatiotemporal Gene Expression Analysis of the Caenorhabditis elegans Germline Uncovers a Syncytial Expression Switch. Genetics. 2018;210(2):587–605.

15. 15. Cadena Del Castillo CE, Hannich JT, Kaech A, Chiyoda H, Brewer J, Fukuyama M, et al. Patched regulates lipid homeostasis by controlling cellular cholesterol levels. Nature communications. 2021;12(1):4898.

16. Kuwabara PE, Lee MH, Schedl T, Jefferis GS. A C. elegans patched gene, ptc-1, functions in germ-line cytokinesis. Genes Dev. 2000;14(15):1933–44.

17. Joshi PM, Riddle MR, Djabrayan NJ, Rothman JH. Caenorhabditis elegans as a model for stem cell biology. Dev Dyn. 2010;239(5):1539–54.

18. Hubbard EJ. Caenorhabditis elegans germ line: a model for stem cell biology. Dev Dyn. 2007;236(12):3343–57.

19. Tolkin T, Mohammad A, Starich TA, Nguyen KCQ, Hall DH, Schedl T, et al. Innexin function dictates the spatial relationship between distal somatic cells in the Caenorhabditis elegans gonad without impacting the germline stem cell pool. Elife. 2022;11.

20. Kimble J, Crittenden SL. Germline proliferation and its control. WormBook. 2005:1–14.

21. Cao W, Fan Q, Amparado G, Begic D, Godini R, Gopal S, et al. A nucleic acid binding protein map of germline regulation in Caenorhabditis elegans. Nature communications. 2024;15(1):6884.

22. Gopal S, Amran A, Elton A, Ng L, Pocock R. A somatic proteoglycan controls Notch-directed germ cell fate. Nature communications. 2021;12(1):6708.

23. Hall BA, Piterman N, Hajnal A, Fisher J. Emergent stem cell homeostasis in the C. elegans germline is revealed by hybrid modeling. Biophys J. 2015;109(2):428–38.

24. Amran A, Pigatto L, Farley J, Godini R, Pocock R, Gopal S. The matrisome landscape controlling in vivo germ cell fates. Nature communications. 2024;15(1):4200.

25. Kimble J, Crittenden SL. Controls of germline stem cells, entry into meiosis, and the sperm/oocyte decision in Caenorhabditis elegans. Annu Rev Cell Dev Biol. 2007;23:405–33.

26. Crittenden SL, Bernstein DS, Bachorik JL, Thompson BE, Gallegos M, Petcherski AG, et al. A conserved RNA-binding protein controls germline stem cells in Caenorhabditis elegans. Nature. 2002;417(6889):660-3.

27. Seidel HS, Smith TA, Evans JK, Stamper JQ, Mast TG, Kimble J. C. elegans germ cells divide and differentiate in a folded tissue. Dev Biol. 2018;442(1):173–87.

28. Gopal S, Boag P, Pocock R. Automated three-dimensional reconstruction of the Caenorhabditis elegans germline. Dev Biol. 2017;432(2):222–8.

29. Seidel HS, Kimble J. Cell-cycle quiescence maintains Caenorhabditis elegans germline stem cells independent of GLP-1/Notch. Elife. 2015;4.

30. Gibieza P, Petrikaite V. The regulation of actin dynamics during cell division and malignancy. Am J Cancer Res. 2021;11(9):4050–69.

31. Cao M, Zou X, Li C, Lin Z, Wang N, Zou Z, et al. An actin filament branching surveillance system regulates cell cycle progression, cytokinesis and primary ciliogenesis. Nature communications. 2023;14(1):1687.

32. Bidet M, Joubert O, Lacombe B, Ciantar M, Nehme R, Mollat P, et al. The hedgehog receptor patched is involved in cholesterol transport. PLoS One. 2011;6(9):e23834.

33. Shamsuzzama, Lebedev R, Trabelcy B, Langier Goncalves I, Gerchman Y, Sapir A. Metabolic Reconfiguration in C. elegans Suggests a Pathway for Widespread Sterol Auxotrophy in the Animal Kingdom. Current biology : CB. 2020;30(15):3031–8 e7.

34. Qu Z, Ji S, Zheng S. Glucose and cholesterol induce abnormal cell divisions via DAF-12 and MPK-1 in C. elegans. Aging (Albany NY). 2020;12(16):16255–69.

35. Lasuncion MA, Martinez-Botas J, Martin-Sanchez C, Busto R, Gomez-Coronado D. Cell cycle dependence on the mevalonate pathway: Role of cholesterol and non-sterol isoprenoids. Biochem Pharmacol. 2022;196:114623.

