## Supplemental figures for "Patched regulates cell cycle and tissue architecture in *C. elegans* gonad"

---

### Supplementary Figures

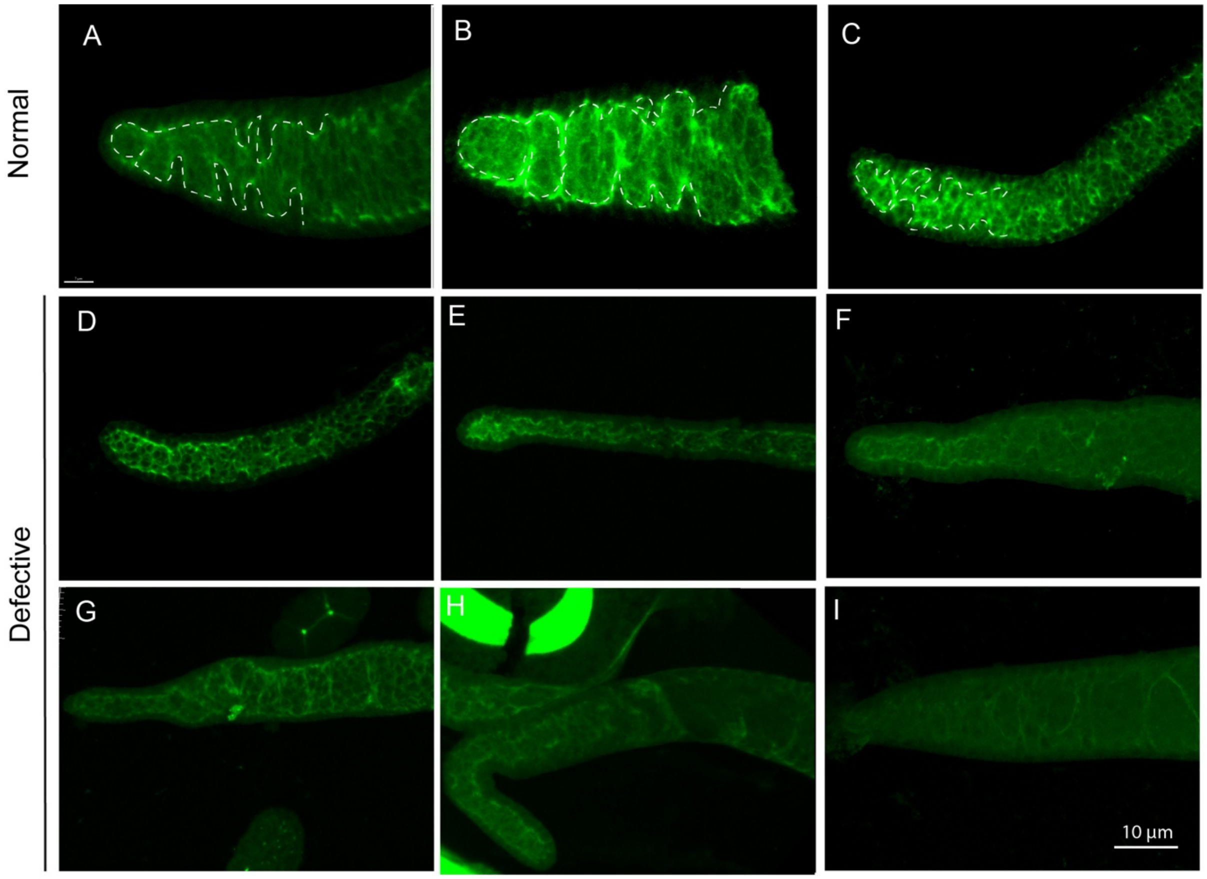


Supplementary figure S1

**Supplementary figure S1. Cytoskeletal morphology of gonad at the distal region.** A-C) Actin cytoskeleton assumes a typical spiral morphology (marked by dotted white lines) at the distal regions of the gonad D-I) Examples of actin cytoskeletal defects observed. Scale bar, 10 µm.

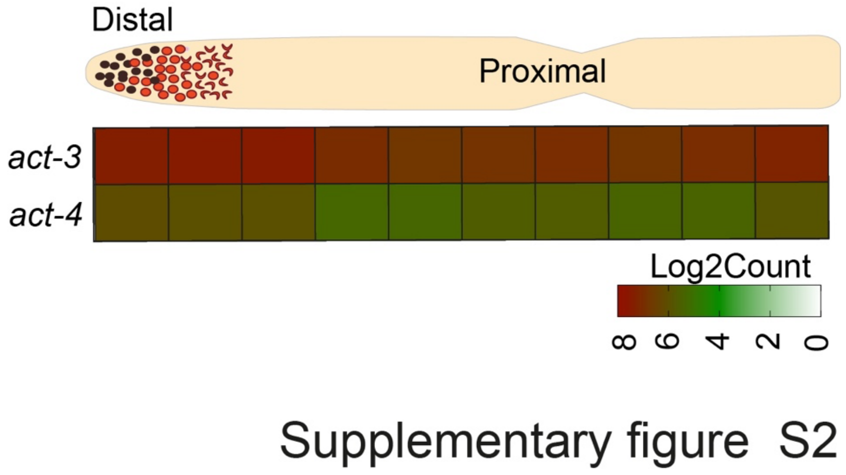


**Supplementary figure S2. Expression of *act-3* and *act-4* in the gonad.** *act-3* and *act-4* expressed throughout the gonad at varying levels from distal to proximal[14].

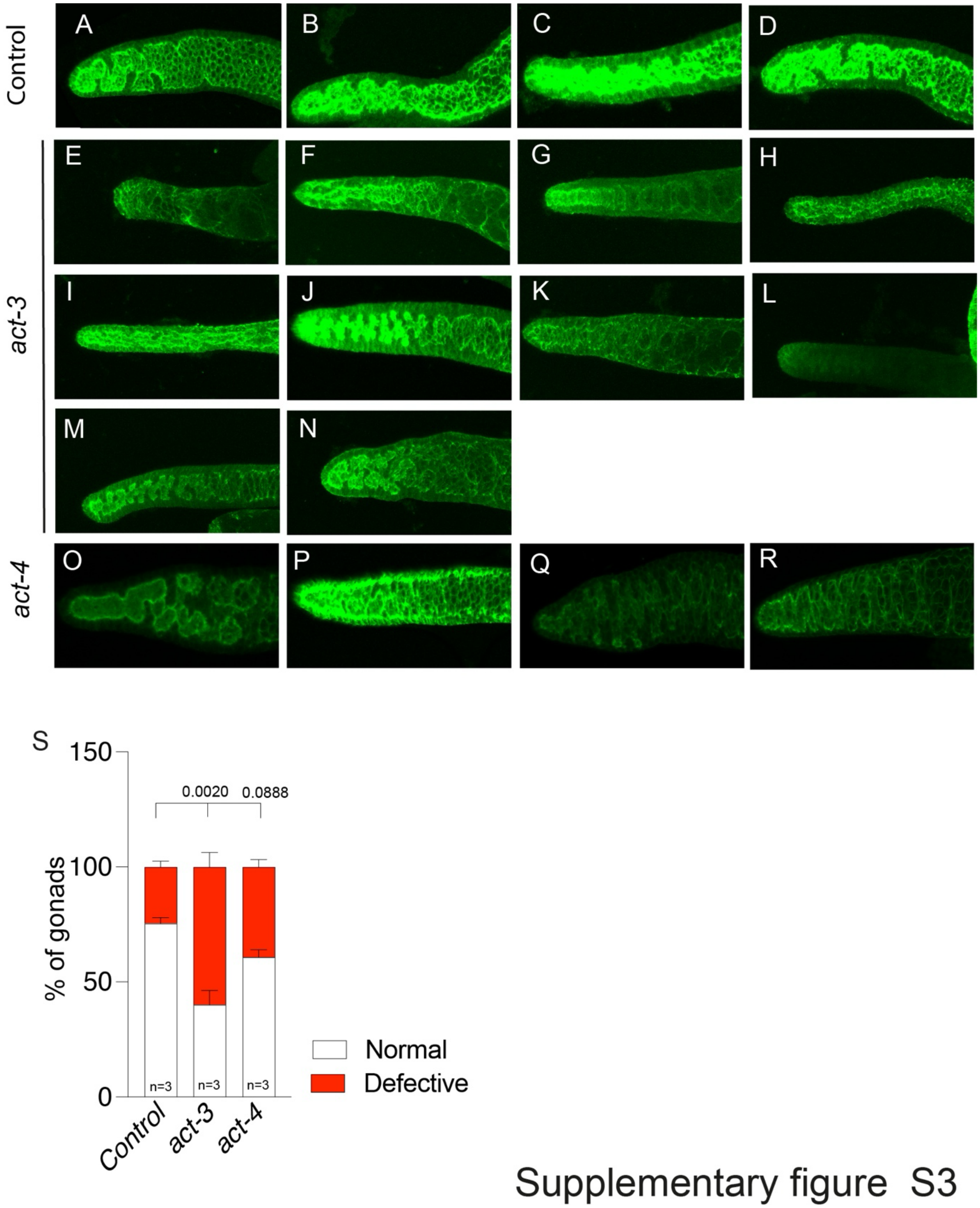


**Supplementary figure S3. Silencing *act-3* and *act-4 i*mpacted gonadal actin**. A-D) Actin phenotype in control animals. E-N) Observed deviations in gonadal actin phenotype after *act-3* RNAi. O-R) Observed deviations in gonadal actin phenotype after *act-4* RNAi. S) Quantification of percentage of gonads with normal and defective cytoskeleton in control and *act-3* or *act-4* silenced animals. Experiment was performed in 3 replicates with a total of 25 germlines for control, 36 germlines for *act-3* RNAi and 20 germlines for *act-4* RNAi. P values were assessed by one-way analysis of variance (ANOVA) with Tukey's multiple comparison test. Error bars indicate SEM. P value < 0.05 = statistically significant. P value < 0.05 = statistically significant.

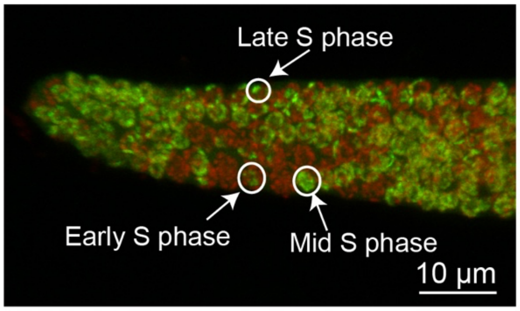


Supplementary figure S4

**Supplementary figure S4. S phase stage identification by EdU staining.** Low EdU intensity = early S phase. Higher EdU intensity = mid S phase. High intensity and punctate staining = late S phase. Scale bar, 10 µm.

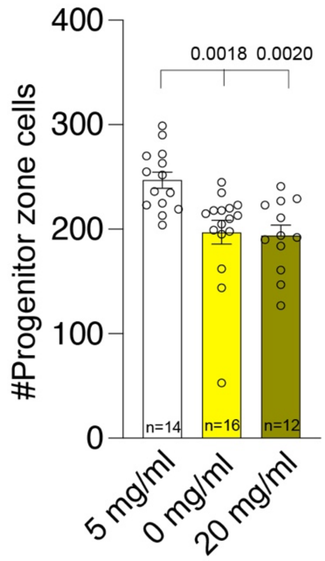


Supplementary figure S5

**Supplementary figure S5. Cholesterol levels regulate cell number in the progenitor zone.** High (20 mg/ml) and no cholesterol (0 mg/ml) conditions negatively affect the germ cell in the progenitor zone compared to normal cholesterol (5mg/ml) levels. P values were assessed by one-way analysis of variance (ANOVA) with Tukey's multiple comparison test. Error bars indicate SEM. P value < 0.05 = statistically significant.

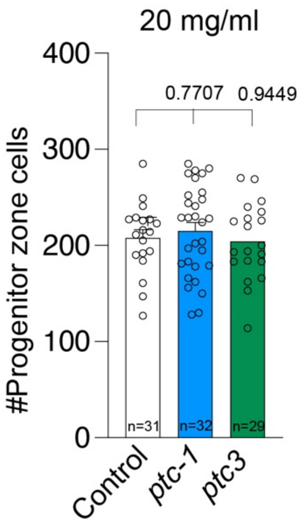


Supplementary figure S6

**Supplementary figure S6. Silencing Patched under high cholesterol conditions did not affect the cell number in the progenitor zone.** Quantification of number of cells in the progenitor zone after silencing *ptc-1* and *ptc-3* under high cholesterol conditions. P values were assessed by one-way analysis of variance (ANOVA) with Tukey's multiple comparison test. Error bars indicate SEM. P value < 0.05 = statistically significant.
